# Autoimmune CD4^+^T cells Cause Meibomian Gland Dysfunction

**DOI:** 10.64898/2026.01.08.697553

**Authors:** Kaitlin K. Scholand, Paola A. Guevara Montoya, Emre Aksan, Yangluowa Qu, Sean Paiboonfungfuang, Laura Schaefer, Elizaveta A. Demianova, Elaine Vo, Sudhir Verma, Elle Joy J San Juan, Zhiyuan Yu, Tarsis F. Gesteira, Vivien J. Coulson-Thomas, Cintia S. de Paiva

**Affiliations:** Department of Ophthalmology, Baylor College of Medicine, Houston, Texas, USA; Department of Biosciences, Rice University, Houston, Texas, USA; College of Optometry, University of Houston, Houston, Texas, USA; Department of Microbiology and Virology, Baylor College of Medicine, Houston, Texas, USA

**Keywords:** Autoimmune, MGD, CD4+T cells, Tregs, dry eye disease

## Abstract

Sjögren disease (SjD) is an autoimmune disease driven by autoreactive CD4^+^T cells that leads to an immune-mediated loss of lacrimal glands. Meibomian glands are lipid-producing glands in the eyelids that help prevent tear evaporation. While the role of T cells in lacrimal gland-mediated destruction is well established, it is unknown whether pathogenic T cells can cause MG dysfunction (MGD). Herein, we investigated whether autoreactive CD4^+^T cells induce MGD and characterized the pathophysiologic mechanisms using an adoptive transfer model. T cells were isolated from CD25KO (CD4^KO^) or wild-type (CD4^WT^) mice, transferred into *Rag1*KO mice. Further, CD4^KO^ cells were co-adoptively transferred with WT regulatory T cells (CD4^KO^+Tregs^WT^). Our results demonstrate that CD4^KO^ recipients had MG dropout, CD4^+^IFN-γ^+^ infiltration, increased MHC II presentation within the periglandular area, MG fibrosis, and decreased lipid production and upregulation of pathways related to inflammation, including Type II interferon signaling. *Rag1*KO, CD4^WT^, and CD4^KO^+Tregs^WT^ recipients exhibited minimal inflammation in the periglandular MG area. These results indicate that autoimmune CD4^+^T cells are sufficient to cause MGD, and healthy young regulatory T cells can prevent T-cell-mediated damage. Taken together, our findings provide mechanistic insights into the pathogenesis autoimmune SjD, and could impact how patients are managed in the clinic.

## Introduction

Dry eye disease (DED) is a chronic condition in which eyes do not get adequate lubrication. It affects millions of people worldwide, with approximately 21 million individuals currently affected in the US(1). DED symptoms vary from mild irritation to severe ocular pain. In more severe cases, the level of pain has been compared to that experienced during angina (2, 3). DED can be caused by increased tear evaporation or by insufficient tear production(2, 3), and, as such, it has been historically divided into “evaporative dry eye “(which includes Meibomian Gland Dysfunction [MGD]) and “aqueous-tear-deficient dry eye” (ADDE), but hybrid or mixed cases exist(4). MGD is the most prevalent cause of DED(5, 6), with clinical studies suggesting that 85% of all DED cases are caused by some form of MGD(7).

Sjögren disease (SjD) leads to a severe form of ADDE and is caused by autoimmune-mediated damage to the lacrimal glands (LGs). SjD is a prototype autoimmune disease that affects the cornea, conjunctiva, and tear fluid-producing LGs. Pathological findings include immune infiltration of T and B cells into salivary gland, LGs and conjunctiva, activation of innate and adaptive immunity, loss of corneal nerves and corneal epitheliopathy(8–13). There is a high frequency of DED patients who present both MGD and SjD-lacrimal gland disease(14–18). However, whether the MGD present in SjD patients is caused by autoimmune-mediated damage remains to be established.

MGs are specialized sebaceous glands within the eyelid’s tarsal plate that secrete lipids (meibum) onto the ocular surface(19). Changes in the quality and quantity of meibum affect its ability to stabilize the tear film, causing tear film evaporation(1). Although the exact etiology of MGD remains unknown, several risk factors have been identified (age, sex, poor diet, diabetes, medical history with certain prior eye surgeries, and autoimmune diseases(20–23)). Conventionally, MGD is believed to be largely caused by the obstruction of MG secretory ducts, leading to the stasis of meibum within the duct and backpressure within the gland, ultimately triggering MG atrophy(19). However, a decrease in cell proliferation within the basal layer of the MG has been observed with aging, reducing the number of cells entering meibocyte differentiation necessary for holocrine secretion and leading to MG atrophy and age-related MGD(24). More recently, studies have suggested that a loss in the number of MG progenitor cells and changes in the extracellular matrix with aging also contribute to age-related MGD(24–26).

Evidence exists to suggest that, as with the lacrimal gland, MGs can suffer inflammatory cell infiltration that contributes to MGD(27, 28). Our group has previously shown that alterations within the MG are observed in the desiccating stress model that is widely used as a model of ADDE(29). Moreover, mice lacking the autoimmune regulator AIRE gene, which exhibit autoimmune phenotypes resembling those observed in SjD, have increased numbers of CD45^+^, CD11b^+^, and CD4^+^ immune cells in and around the MG(28). Moreover, CD45^+^ cells, increased expression of inflammatory markers, and increased fibrosis have been identified in the periglandular area of the MG in Cu, Zn-Superoxide Dismutase-1 (Sod1) KO mice when compared to wild-type (WT) mice(30). Likewise, IL-17^+^CD4^+^T cell infiltration causes MG plugging, fibrosis, and neutrophil recruitment in a chronic model of allergic eye disease(31, 32). In two different models of ocular graft-versus-host-disease, a bone marrow transplant with T cells was sufficient to drive MGD(33, 34). Finally, meibum with reduced quality has been correlated with CD45^+^ cells infiltrating the MG acini and/or ducts(35). Thus, prior work suggest MGs are susceptible to immune cell-mediated inflammation; however, evidence that inflammatory cell infiltration directly causes MGD is still lacking.

This study investigated whether autoimmune CD4+ T cells infiltrate the MG and trigger an inflammatory response leading to MG atrophy and contributing to MGD, as observed in the LGs.

## Material and Methods

### Study approval

The Institutional Animal Care and Use Committee at Baylor College of Medicine approved all animal experiments (AN-6491) and all studies adhered to the Association for Research in Vision and Ophthalmology for the Use of Animals in Ophthalmic and Vision Research and the NIH Guide for the Care and Use of Laboratory Animals(36). All animal experiments were conducted at the Ocular Surface Center, Department of Ophthalmology, Baylor College of Medicine, Houston, Texas. Samples were collected and transported to the College of Optometry, Houston, Texas for analysis.

### Animals and Sex as a Biological Variable

Heterozygous breeder pairs of *Il2ra*^+/-^ mice on a C57BL/6 background *(Il2ra* or CD25, B6.129S4-Il2ra^tm1Dw^/J, stock 002952) and breeder pairs of *Rag1*^+/-^ mice (Recombination activating gene 1; B6.129S7-Rag1^tm1Mom^/J, stock :002216) were purchased from Jackson Laboratories (Bar Harbor, Maine) for establishing breeder colonies. The genotype of the CD25KO mice was confirmed according to the Jackson Laboratories’ protocol using a commercial vendor (Transnetx, Cordova, Tennessee). Young C57BL/6J donor mice were purchased from Jackson Laboratories (stock 000664, Bar Harbor, Maine) and were used between 8-10 weeks of age. Mice were housed in specific pathogen-free facilities at the Baylor College of Medicine on diurnal cycles of 12 hours/light and 12 hours/dark with *ad libitum* access to food and water and environmental enrichment.

*Il2ra*^-/-^ mice (CD25KO, n = 56) and *Il2ra*^+/+^ mice (CD25^+/+^, WT, n = 106) were used at 8 weeks of age at the start of the experiment. *Rag1*KO mice aged 8 weeks to 5 months were used (n = 130, 80 males and 50 females). Experiments were performed with both male and female animals, treating sex as a biological variable. Some endpoints were performed preferentially with male mice due to their greater availability and the more pronounced phenotype.

### CD4^+^ T cell Isolation, Adoptive Transfer and Co-Adoptive Transfer

Single-cell suspensions were prepared from the spleens and draining lymph nodes of either CD25^+/+^ (wild-type) or CD25KO mice. CD4^+^ T cells were isolated by negative selection using the CD4^+^ T Cell Isolation Kit (Miltenyi Biotec, Bergisch Gladbach, Germany) following the manufacturer’s instructions. Two million CD4^+^T cells were injected intraperitoneally into sex-matched *Rag1*KO mice (CD4^WT^ and CD4^KO^, respectively). Co-adoptive transfer experiments used 2×10^6^ CD4^+^T cells isolated from the CD25KO with 200,000 Tregs (CD4^+^GITR^+^CD25^+^ cells) sorted from wild-type mice, as in(37, 38).

Mice were euthanized at five weeks after adoptive transfer. Tarsal plates were processed and imaged as described below (n = 30-43 biological replicates/group) and then harvested and used for histology/immunohistochemistry/Oil Red O (ORO, at least n = 6 biological replicates/group), flow cytometry (n = 11-13 biological replicates/group), bulk RNA sequencing (n = 5 biological replicates/group), and qPCR validation (n = 6-11 biological replicates/group). Efforts were made to use the same mice across multiple endpoints to reduce sample size whenever possible. The final sample size per endpoint is shown in the figure legends.

### Collection and Processing of Eyelids

Mice were euthanized by isofluorane inhalation in a glass bell-jar, followed by cervical dislocation. Immediately after euthanasia, a bright-field image was acquired using a Nikon LMZ1500 stereoscope, with the FS-Di1 camera (Nikon, Melville, NY). Eyelids were dissected and either photographed or fixed in 4% buffered paraformaldehyde or tarsal plate dissection, which were either lysed in RNA lysis buffer and stored at -80 °C for mRNA isolation or processed for flow cytometry. For whole mount imaging, eyelids were trimmed under a dissecting microscope (Nikon LMZ1500, Camera FS-Di1) or (Leica S9E, Leica Microsystems Inc., IL, USA) and excess tissue was dissected to evidence the MGs and the eyelids mounted as flat mounts between two glass slides and imaged under a stereomicroscope using a Discovery.V12 (ZEISS, Oberkochen, Germany) fluorescence stereomicroscope or a Nikon LMZ1500 using white light and the 488 channel, as previously shown(25, 26). All samples included in this study were first analyzed by flat mount prior to processing. Fixed tissues were stored at 4C until further analysis.

### Measurement of MG area, Number of Glands and Enlarged Ducts

The outer perimeter of the MGs was demarcated using the curvature pen tool in Adobe Photoshop (San Jose, California). The MG area was quantified using ilastik (https://www.ilastik.org/) and ImageJ. MG area calculations were either for both superior and inferior eyelids, referred to as total MG area, or only of the inferior eyelid, referred to as inferior MG area. Since the MGs in the nasal region of the eyelid are sometimes inadvertently removed during flat mount dissections, we opted to exclude these glands from all area quantification analysis. The area measurements obtained for both the right and left eyelids were averaged for each mouse and presented as a single data point on the graph. Further, the total number of glands within the superior and inferior eyelids were manually counted in digital images of eyelid flat mounts by two independent investigators in a blinded manner. Glands were included in the total count only if they retained an identifiable glandular structure. Areas of MG dropout, defined as total loss of acinar tissue, were excluded from the final count. Enlarged ducts were also manually counted in digital images of flat mounted eyelids. For such, the brightness of the bright field images was increased beyond the level of saturation for the MGs (an additional 100 gray levels in Photoshop) to facilitate the visualization of the dilated collecting ducts. The final sample size for each experimental group is provided in the figure legends.

### Immunohistochemistry

Eyelids were flat-mounted and embedded in Tissue-Tek embedding medium (Sakura Finetek USA, Inc., Torrance, Callifornia) and 10 µm coronal cryosections obtained through the eyelids using a Leica CM 1950 (Leica, Buffalo Grove, IL, USA) cryostat and collected as sections on Fisherbrand SuperfrostPlus Gold microscope slides (Thermo Scientific, Wilmington, Delaware). Upon use, slides were placed on a slide warmer (XH-2002; Premiere, C&A Scientific, Sterling, Virginia) set at 60°C for 30 minutes and washed with PBS. Tissues were blocked with 10% fetal bovine serum (FBS, Thermofisher Scientific) prepared in PBS containing 0.01 M saponin (ThermoFisher Scientific). Sections were incubated with primary antibodies anti-Keratin 14 (PRB-155P; Covance, Princeton, NJ), anti-A-I/A-E (558996, BD Biosciences, Milpitas, California), anti-MHC class II (PA-14-5321-82, Invitrogen, Eugene, Oregon), anti-LAMP3 (ThermoFisher-MA1-19650, ThermoFisher Scientific, Rockford, Illinois), anti-α smooth muscle actin (14-9760-82, clone 1A4, Millipore Sigma, Burlington, Massachusetts), and biotinylated HA binding protein (HABP, 385,911, Millipore Sigma, Burlington, Massachusetts). Sections were left for one hour at room temperature. Secondary controls were carried out in parallel by omitting the primary antibody and did not yield any specific staining (results not shown). Some sections were further stained with Alexa-647 Phalloidin (Invitrogen) for one hour at room temperature and nuclei counter stained with DAPI. Slides were mounted in FluormountG® (Invitrogen) and imaged under a LSM 800 confocal microscope (ZEISS).

Alternatively, the eyes and ocular adnexa were excised, snap-frozen in liquid nitrogen in an OCT-filled plastic cassette and stored at -80°C until use. 5-µm sagittal cryosections were cut using a cryostat (HM525NX, Epredia, Kalamazoo, Michigan). Sections were thawed, fixed in cold acetone, washed in PBS and stained with anti-CD4 (553647, BD Biosciences) and anti-Foxp3 (14-5773-82, ThermoFisher) antibodies at room temperature for one hour. After washing three times, sections were incubated with the appropriate biotinylated secondary antibody (559286, BD Biosciences) and a Vectastain Elite kit (PK-7100, Vector Laboratories, Burlingame, California). Color development was performed using NovaRed reagent (SK-4800, Vector Laboratories), as previously described(39).

### Oil Red O staining

Eyelids were immersed in freshly prepared Oil-Red-O solution (Sigma-Aldrich, St. Louis, MO) and maintained at room temperature under gentle agitation for two hours, protected from light. The eyelids were then de-stained with deionized (DI) water, flat mounted in FluormountG® (Invitrogen) and imaged under a stereomicroscope (Discovery.V12, ZEISS, Oberkochen, Germany) using the 555 channel.

### Masson’s Trichrome staining

Coronal MG cryosections were stained with Trichrome stain (Gomori One-Step Light Green, Newcomer Supply Inc., Waunakee, Wisconsin) following the manufacturer’s instructions. In short, frozen sections were stained with Hematoxylin (Harris Modified, Newcomer Supply Inc.,), rinsed in running water and incubated in Trichrome stain at 40 °C for 20 minutes. Subsequently, the slides were differentiated in 0.25% acetic acid, after which they were dehydrated through 95% and 100% ethanol washes. The sections were then washed with three xylene washes and mounted using Permount® (Fisher Scientific). Spiral images were captured under a Leica DMi8 inverted microscope.

### Flow cytometry

The superior and inferior tarsal plates from both right and the left eyelids were pooled and placed in a solution of 0.1% collagenase D (from Clostridium histolyticum, SigmaAldrich, St. Louis, Missouri) in Hank’s Balanced Salt Solution (Gibco, ThermoFisher), minced, and incubated at 37°C for one hour. In the last fifteen minutes, DNAse I (MilliporeSigma) was added to each suspension. In the last five minutes, 0.05% EDTA-Tripsin (Gibco, ThermoFisher) was added. Suspensions were neutralized with RPMI (Gibco, ThermoFisher) and filtered through 100µm CellTrics filters (Sysmex, Kobe, Japan) before staining. Suspensions were stimulated with ionomycin (1ug/ul, SigmaAldrich) and PMA (1ug, SigmaAldrich) for five hours in an incubator set at 37°C and 5% CO_2_. In the last 2 hours, Golgi Stop (1 μL/suspension; BD Biosciences) was added. Single-cell suspensions were incubated with CD16/32 (clone 2.4G2; BD Biosciences) to block Fc receptors for ten minutes on ice before being stained with an infrared fluorescent reactive live/dead dye diluted 1:3000 (Invitrogen, ThermoFisher) for fifteen minutes.

Cells were immediately fixed and permeabilized with the BD Pharmingen Transcription Buffer set (BD Biosciences) following the manufacturer’s protocol. Cells were then stained overnight at 4°C with the following: CD45-BV510 (clone 30-F11; BioLegend, San Diego, California), CD3-BB700 (clone 145-2C11, BD Biosciences), CD4-FITC (clone RM4-5; Invitrogen ThermoFisher), FoxP3-APC (clone FJK-16s, BioLegend), IL-17-PE (clone eBio1787; Invitrogen ThermoFisher), and IFN-γ-Pacific Blue (clone XMG1.2, BioLegend). The next day, suspensions were washed four times before data acquisition on a BD Canto II Benchtop cytometer with BD Diva software version 6.7 (BD Biosciences). Final data were analyzed using FlowJo software version 10 (Tree Star, Ashland, Oregon). The following gating strategy was used for these experiments: cells were identified by forward scatter area versus side scatter area; doublets were discriminated by forward scatter height versus forward scatter area (singlet 1), followed by side scatter height versus side scatter area (singlet 2); thereafter, dead cells were excluded by gating live fixable infrared dye versus side scatter; and subsequently, CD45^+^ cells were gated and then CD4^+^ were selected to calculate the frequency of intracellular markers IFN-γ^+^, IL-17^+^, or FoxP3^+^ as described in each figure.

### RNA Isolation and Bulk RNA Sequencing

Tarsal plates from *Rag1*KO, CD4^WT^, CD4^KO^, and CD4^KO +^Tregs^WT^ mice were excised, and added to an Eppendorf tube containing RNA lysis buffer (Qiagen, Hilden, Germany), snap-frozen in liquid nitrogen and stored at - 80 °C. Total RNA was extracted using a QIAGEN RNeasy Plus Micro RNA isolation kit (Qiagen) following the manufacturer’s protocol. A double-stranded DNA library was created at Baylor College of Medicine’s Genomics and RNA Profiling Core. Briefly, using 100ng of total RNA (measured by pico green), an oligodT primer containing an Illumina-compatible sequence at its 5’ end was hybridized to the RNA, and reverse transcription was performed using a Lexogen kit. Second-strand synthesis was initiated by a random primer containing an Illumina-compatible linker sequence at its 5’ end. The purified double-stranded library was then amplified and purified. The resulting libraries were quantitated using Qubit 2.0 (ThermoFisher) and fragment size was assessed with the Agilent Bioanalyzer (Agilent Technologies, Santa Clara, California). A qPCR quantitation was performed on the libraries to determine the concentration of adapter-ligated fragments using Applied Biosystems ViiA7 ^TM^ Real-Time PCR System and a KAPA Library Quant Kit (Waltham, Massachusetts). All samples were pooled equimolarly, re-quantitated by qPCR, and reassessed on the bioanalyzer. Using the concentration from the ViiA7 ^TM^ qPCR machine above, 1.8pM of the equimolarly pooled library was loaded onto NextSeq 500 high-output flow cell (Illumina, San Diego, California). PhiX Control v3 adapter-ligated library (Illumina) was spiked in at 1% by weight to ensure balanced diversity and to monitor clustering and sequencing performance. A single-read 75-base pair cycle run was used to sequence the flow cell. An average of 21 million reads per sample was sequenced. The FastQ file generation was executed using Illumina’s cloud-based informatics platform, BaseSpace Sequencing Hub (Illumina).

Data were analyzed by ROSALIND® (https://rosalind.bio/), with a HyperScale architecture developed by ROSALIND, Inc. (San Diego, California) as previously described(40–42). Enrichment was calculated relative to a set of background genes relevant to the experiment.

### Quantitative PCR

RNA concentration was measured, and cDNA was synthesized using the Ready-To-GoTM You-Prime FirstStrand kit (Cat# 27926401, GE Healthcare, Chicago, Illinois). Quantitative real-time PCR was performed with specific minor groove binder probes for CXCL9 (*Cxcl9*, Mm00434946_m1), CXCL10 (*Cxcl10*, Mm00445235_m1), cathepsin S (*Ctss*, Mm01255859_m1), mannose receptor C-type 1 (*Mrc1*, Mm00485148_m1), Col6a5 (*Col6a5*, Mm01231908_m1), Col6a6 (*Col6a6*, Mm00556810_m1), TNF (*Tnf*, Mm00443258_m1), IL-1β (*Il1b*, Mm00434228_m1), interferon-γ (*Ifng*, Mm01168134_m1), class II transactivator (*Ciita*, Mm00482914_m1), and hypoxanthine–guanine phosphoribosyltransferase 1 (*Hprt1*, Mm00446968_m1). Two different housekeeping genes were tested (GAPDH and HPRT1). Since we found both to be equivalent and there was no difference among the groups, the HPRT1 gene was used as an endogenous reference for each reaction. The results of real-time PCR were analyzed using the comparative delta deltaCT method, and the data were normalized to the HPRT1 cycle threshold value. Specific outliers were identified using GraphPad software.

### Statistical analysis

The sample size was calculated with StatMate2 Software (GraphPad Software, San Diego, California) based on pilot studies. Statistical analyses were performed with GraphPad Prism software (version 10.6.1, GraphPad). Data were first evaluated for normality with the Kolmogorov-Smirnov normality test. Appropriate parametric (t-test) or non-parametric (Mann-Whitney) statistical tests were used to compare the groups. Whenever adequate, one-way or two-way ANOVA or Kruskal-Wallis followed by post hoc tests were used. The final sample per experiment is shown in the legends

### Data availability

Bulk RNA sequencing data can be found at GSE314353. All other data will be provided by the authors upon request.

## Results

### Autoimmune CD4^+^T cells within the MG and tarsal plate cause MGD

We and others have shown that CD25 (*Il2ra*) knock-out (CD25KO) mice, a mouse model of SjD, develop systemic autoimmunity, and severe SjD-like ocular and lacrimal gland alterations linked to overly activated T cells and loss of regulatory T cells (Tregs)(39, 43–48). Previously, we used this model to characterize pathogenic autoreactive CD4^+^ T cell-mediated destruction of the tear-secreting LGs(38, 39). We used the same adoptive transfer model (“→”) to verify whether CD25KO CD4^+^T cells cause MGD. We isolated CD4^+^ T cells from ocular draining lymph nodes and spleens of 8-week-old CD25KO mice using magnetic beads (CD4^KO^, >95% purity, Miltenyi) and adoptively transferred them intraperitoneally into age- and sex-matched *Rag1*KO mice. As a control, CD4^+^ T cells were isolated from ocular draining lymph nodes and spleens from 8-week-old CD25^+/+^ wild-type littermates and adoptively transferred into age and sex-matched *Rag1*KO mice (CD4^WT^) (Supplemental Figure 1A). Age-matched *Rag1*KO mice were included as naïve controls. Mice were euthanized 5 weeks post-adoptive transfer, at which point severe lacrimal gland disease was reported(38, 39). At euthanasia, the ocular surface and eyelids of CD4^WT^ appeared normal, comparable to those of the naïve *Rag1*KO mice, while some CD4^KO^ eyes had periocular edema and a small palpebral aperture (data not shown). The eyelids were excised, mounted for whole-mount analysis, and imaged using a stereomicroscope (Figure 1B,C). Minimal MG dropout was observed in the CD4^WT^ group and *Rag1*KO mice, while CD4^KO^ recipients presented MG atrophy, which was more significant in the lower lid (Figure 1D). The total MG area and the lower lid MG area were calculated. CD4^KO^ mice presented a significant decrease in the MG area within the lower eyelid when compared to *Rag1*KO and CD4^WT^ mice (Figure 1B-D). CD4^KO^ mice presented MG atrophy occurring randomly along the length of the gland(Figure 1B-D), rather than primarily in the region most distal to the eyelid margin, as previously observed with age-related MGD(25). The number of MGs and the number of enlarged collecting ducts were counted in the superior and inferior eyelids of *Rag1*KO, CD4^WT^, and CD4^KO^ mice, and a significant decrease in the total number of MGs and a significant increase in the number of enlarged ducts was observed in CD4^KO^ mice when compared to *Rag1*KO and CD4^WT^ mice, with the differences being more pronounced in the inferior lid (Figure 1E and F).

**Figure 1:**
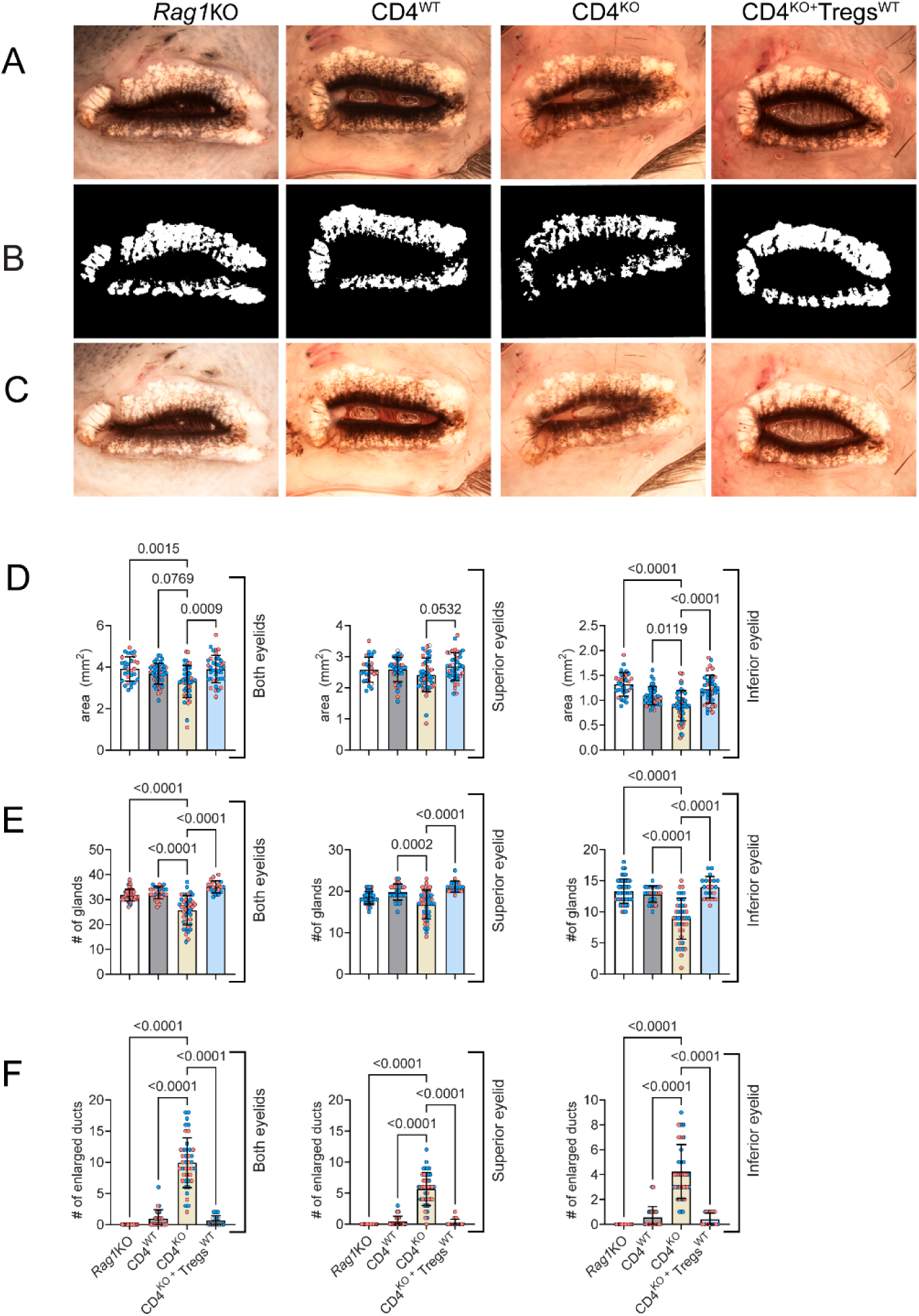
Adoptive transfer of autoreactive CD4^+^ T cells leads to MG atrophy and reduced MG area, MG drop-out and MG plugging. A. Representative digital image of whole-mounted eyelids under a stereomicroscope using white light. **B**. Digital images were analyzed as described in the methods, and the MG area was quantified in the binarized images. **C.** Plugged and enlarged ducts were counted in overexposed digital images to facilitate visualization. **D-F**. Quantifications of MG area (D, n = 30-43/group), number of glands (**E**, n = 11-35), and number of plugging/enlarged ducts (**F**, n = 11-35) in both eyelids, superior and inferior, are presented as graphs with mean ± SD. Each data point on the graph represents a single animal. Male samples are shown as blue dots, and female samples as red dots. One-way ANOVA with Dunn’s multiple comparisons test was used, and the p-values are presented within the graph.

CD25KO mice have impaired Treg function and an excess of autoreactive CD4^+^ T cells(44). To verify whether Tregs from healthy wild-type littermates can prevent the MG atrophy caused by autoreactive T-cells, CD4^KO^ T cells were co-adoptively transferred with WT Tregs at the biological ratio of 10:1 ratio (sorted as CD4^+^GITR^+^CD25^+^ cells from spleens and lymph nodes, as in (37, 38))(CD4^KO^+Treg^WT^). At euthanasia, the ocular surface and eyelids of CD4^KO^ +Treg^WT^ appeared normal, comparable to those of CD4^WT^ and *Rag1*KO mice. CD4^KO^+Treg^WT^ mice presented significantly less MG atrophy compared to CD4^KO^ mice, with total MG area and inferior lid MG area comparable to that of *Rag1*KO and CD4^WT^ mice (Figure 1A, B, D). CD4^KO^+Treg^WT^ mice presented a significantly higher number of MGs and significantly fewer enlarged collecting ducts in the upper and lower eyelids when compared to CD4^KO^ mice, with a comparable number of MGs and enlarged ducts to the *Rag1*KO and CD4^WT^ mice (Figure 1E and F). Therefore, CD4^KO^ recipients presented MG atrophy, drop-out, and plugging (indicated by enlarged ducts), whereas CD4^KO^+Treg^WT^ recipients did not present any signs of MG pathology. This suggests that autoreactive T cells can mediate autoimmune MGD, but regulatory T cells can prevent MGD when co-transferred (**Supplemental Figure 1 and Figure 1).**

### Th1 cells infiltrate the MG after adoptive transfer

To identify potential mechanisms by which T cells cause MG atrophy and dropout, tarsal plates were collected and processed for immunostaining and flow cytometry. First, eyelid cryosections were stained with an anti-CD4 antibody. CD4^+^T cells were identified surrounding MG acini in high abundance in the CD4^KO^ recipient group compared with all other groups. However, CD4^KO^+Treg^WT^ eyelids showed minimal T cell infiltration when compared to the CD4^KO^ group, indicating the co-adoptive transfer of Tregs prevents CD4^+^T cell infiltration (Figure 2A). Moreover, Foxp3^+^ T cells were identified in the periglandular MG area of the CD4^KO^+Treg^WT^ group by immunohistochemistry (Figure 2B).

**Figure 2:**
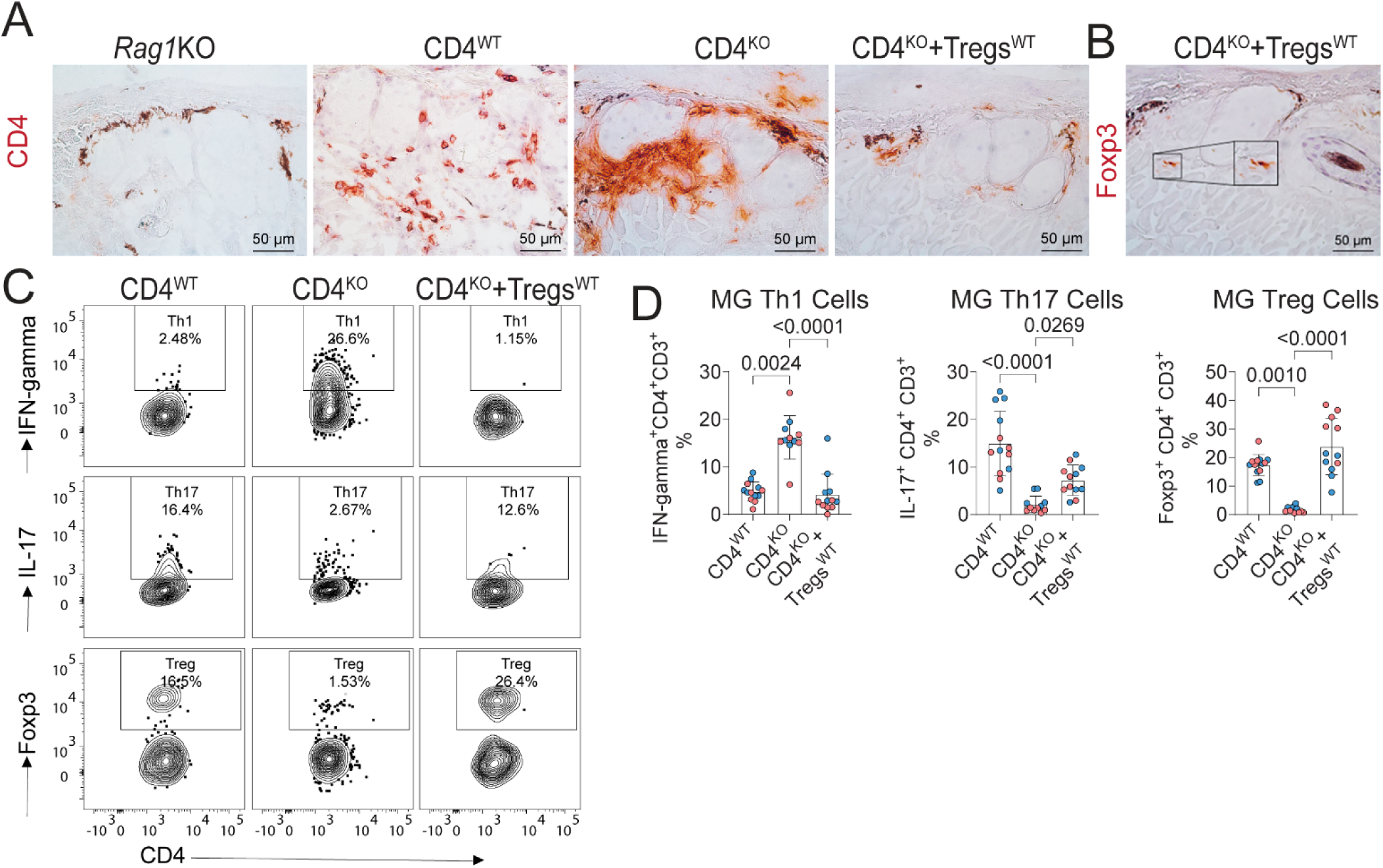
CD4^+^T cells infiltrate MG in *Rag1*KO recipients. Tarsal plates were collected and processed for immunostaining and flow cytometry to assess whether T cells directly attack the MGs. **A-B** Representative images of eyelid cryosections stained with anti-CD4 (A, maroon) or Foxp3 (**B**, maroon) with light hematoxylin counterstaining. Black = melanin. Scale bar = 50 µm. **C**. Flow cytometry of *Rag1*KO tarsal plates identifies differing T cell subsets depending on the adoptive transfer cell source. Representative flow plots of regulatory T cells (CD4^+^Foxp3^+^), Th1 (CD4^+^IFN-γ^+^), and Th17 (CD4^+^IL-17A^+^) cells. **D.** Cumulative data of flow cytometry measurements measuring Th1, Th17 and Treg frequencies in meibomian glands (MG). Each dot represents one mouse, n = 11-13/group; males are presented with blue dots while females are presented with red dots. Mean± SD. One-way ANOVA with Dunn’s multiple comparisons test. P-value as shown.

T cell infiltration into the tarsal plates and MGs was also assessed by flow cytometry. Tarsal plates were prepared as a single-cell suspension and analyzed by flow cytometry. In the CD4^WT^ group, intracellular staining identified a low frequency of Th1 (CD4^+^IFN-γ^+^) and Th17 (CD4^+^IL17^+^) cells and a high frequency of CD4^+^Foxp3^+^ cells (Figures 2C and D). In contrast, CD4^KO^ recipients had a significant increase in Th1 and a significant decrease in Th17 cells (Figure 2C and D). Like the CD4^WT^ mice, CD4^KO^+Treg^WT^ recipients showed a low frequency of Th1 cells and an increase in CD4^+^Foxp3^+^ cells (Figure 2C and 2D). These results suggest that Th1 cells are the main pathogenic T cell subtype that infiltrates the MG of the CD4^KO^ recipients in this model.

### Autoimmune MGD is accompanied by significant transcriptome changes

To further investigate the mechanism by which autoimmune CD4^+^T cells drive MG atrophy, the tarsal plates were then processed and analyzed by bulk RNA sequencing. First, we compared all adoptive transfer groups with naïve *Rag1*KO mice using a 1.5-fold change threshold and a p-adjusted value <0.05, as implemented in ROSALIND (Supplemental Figure 2). There were minimal changes between the CD4^WT^ and *Rag1*KO groups (only 28 differentially expressed genes, [DEGs], Supplemental Figure 2), while the CD4^KO^ MGs displayed the greatest number of DEGs (1153, Supplemental Figure 2B). The CD4^KO^+Treg^WT^/*Rag1*KO showed a ∼10-fold decrease compared to 1153 DEGs from the CD4^KO^/*Rag1*KO (Supplemental Figure 2C). ROSALIND Meta-analysis of these comparisons showed that the upregulated DEGS mapped to BioPlanet Pathways of “Immune system, “immune system signaling by interferons, interleukins, and growth hormones, “T cell receptor regulation of apoptosis,” “IL-2 signaling pathway,” and “Interferon signaling” (Supplemental Figure 2D). Downregulated pathways included “Metabolism”, “Drug Metabolism cytochrome P450“, and “Metapathway biotransformation.”

Since *Rag1*KO mice are devoid of T and B cells, we directly compared gene expression after adoptive transfer between CD4^KO^ and CD4^WT^ MGs, and then CD4^KO^+Treg^WT^ to CD4^KO^, to investigate the effects of pathogenic T cells in the MGs. We observed that 541 DEGs were present in the CD4^KO^ to CD4^WT^ comparison (57 down and 484 upregulated), while 947 DEGs (777 down and 170 up) were present in the CD4^KO^+Treg^WT^/CD4^KO^ comparison (Figure 3 A-B). The volcano plots in Figure 3A and B highlight some DEGs that were upregulated in the CD4^KO^ recipients but were downregulated in the CD4^KO^+Treg^WT^ mice *(Cxcl9, Cxcl10, Ciita, Cd4, Ifng, Ctss)*.

**Figure 3:**
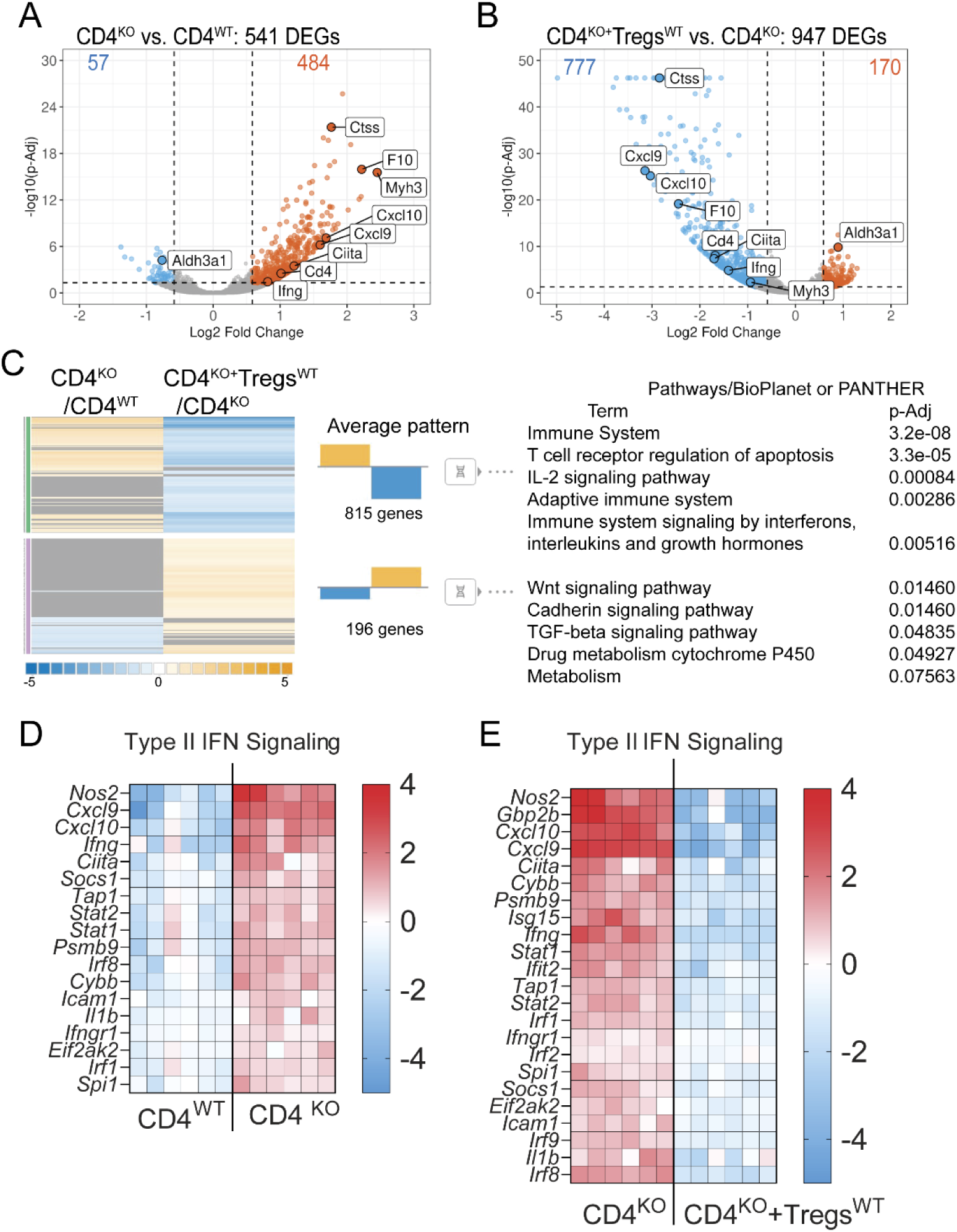
Inflammatory pathways within the MG are differentially modulated in the presence of Tregs. Male tarsal plates (n = 5/group) were collected after 5 weeks post-transfer and lysed. Total RNA was subjected to bulk RNA sequencing. **A-B** Volcano plots displaying the magnitude of change comparing CD4^KO^ to CD4^WT^ transfer (A) and CD4^KO^+Tregs^WT^ to CD4^KO^ **(B)** tarsal plates. **C.** Metanalysis demonstrating unique upregulated (yellow)/downregulated (blue) pathways. **D-E.** Heatmaps of DEGs related to Type II interferon (IFN) signaling in the CD4^KO^ (D) or CD4^KO^+Tregs^WT^ recipients (E).

Meta-analysis of these comparisons revealed that CD4^KO^+Treg^WT^ recipients’ downregulated pathways were related to the “Immune system,” “T cell receptor regulation of apoptosis,” and “IL-2 signaling pathways” among others, while upregulated pathways were related to “Wnt signaling,” “Cadherin signaling,” and the “TGF-β pathway” (Figure 3C). ROSALIND analysis identified that “Type II interferon signaling” was the most upregulated pathway in the CD4^KO^/CD4^WT^ comparison and the most downregulated pathway in the CD4^KO^+Treg^WT^/CD4^KO^ comparison. The supplementary file lists all significant DEGs in these comparisons. Figure 3D and E depict the heatmaps of the DEGS involved in the Type II interferon pathway. The type II Interferon signature gene *Ifng* was upregulated by ∼4-fold in the CD4^KO^ group and downregulated by ∼3-fold in the CD4^KO^+Treg^WT^ group. Other type II interferon DEGs, including IFN-γ-inducible chemokines (*Cxcl9, Cxcl10*) and transcription factors (*Stat1, Stat2*), *Ciita* and interferon-regulated genes (“*irf*”) followed a similar pattern to *Ifng*. These results are consistent with our flow cytometry results, which demonstrated a marked increase in IFN-γ in CD4^KO^ recipients and a decrease in IFN-γ levels when Tregs (CD4^KO^+Treg^WT^) were present.

Next, we validated the bulk RNA sequencing using qPCR in a larger sample size. Consistent with our flow cytometry results, which identified Th1 cells as being pathogenic, qPCR identified that *Ifng* and *Ifng-*related genes (*Ctss, Ciiita, Cxcl9, Cxcl10*) were upregulated in the CD4^KO^ group and downregulated in the CD4^KO^+Treg^WT^ group. *Ctss* encodes cathepsin S, an enzyme that is critical for MHC II presentation*. Ciita* encodes MHC class II transactivator, the master gene regulating MHC II and both genes are upregulated in dry eye(49–52). We also observed a significant increase in the inflammatory transcripts of *Tnf* and *Il1b* in the CD4^KO^ recipient tarsal plates. Expression of these markers was significantly lower in the CD4^KO^+Treg^WT^ group (Figure 4). Taken together, these results indicate profound transcriptome differences are present between the CD4^KO^ to CD4^WT^ groups, and the CD4^KO^ to CD4^KO^+Treg^WT^ comparisons and identified “Type II interferon signaling” as a major signaling pathway.

**Figure 4:**
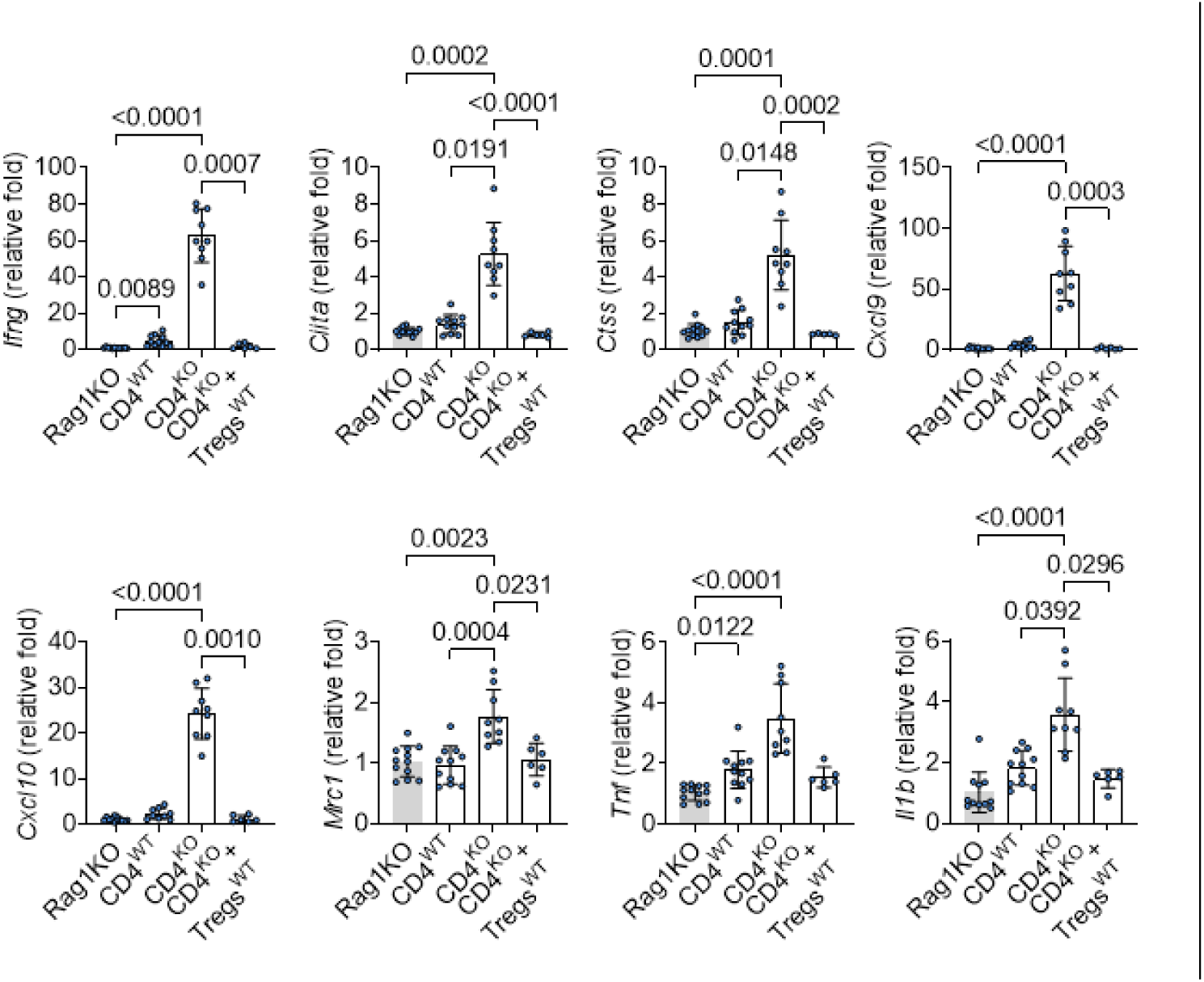
CD4^KO^ recipients have upregulated type II interferon genes. Male tarsal plates were collected and subjected to qPCR-based gene expression analysis, which validated selected DEGs involved in the Th1 response. P-value as shown. Each dot represents one male tarsal plate from separate mice. Mean± SD. One-way ANOVA with Dunn’s multiple comparisons test.

Thereafter, flat-mounted eyelids were cryosectioned, and coronal sections were collected to perform protein validation studies. The histological features of the MG within the coronal sections were compared to the digital images of the whole-mounted eyelids, enabling the identification of atrophic and healthy glands. The outer boundaries of healthy MGs in whole-mount digital images are indicated in green, while those of MGs undergoing atrophy are indicated in orange in Figure 5. *Rag1*KO and CD4^WT^ presented limited MHC class II expression that was restricted to antigen-presenting cells (APCs) (Figure 5A). A significant increase in MHC class II expression was noted surrounding the MGs within the eyelids of CD4^KO^ recipients, with a more pronounced increase in MHC class II staining surrounding MGs undergoing atrophy (Figure 5A). Further, MHC class II expression was not limited to APCs in CD4^KO^ recipients. Co-adoptive transfer of Tregs prevented the increase in MHC class II expression observed in CD4^KO^ mice. Thus, CD4^KO^ MGs have high MHC class II expression by non-immune resident cells, which is prevented by the co-adoptive transfer of T cells in CD4^KO^+Treg^WT^ mice.

**Figure 5:**
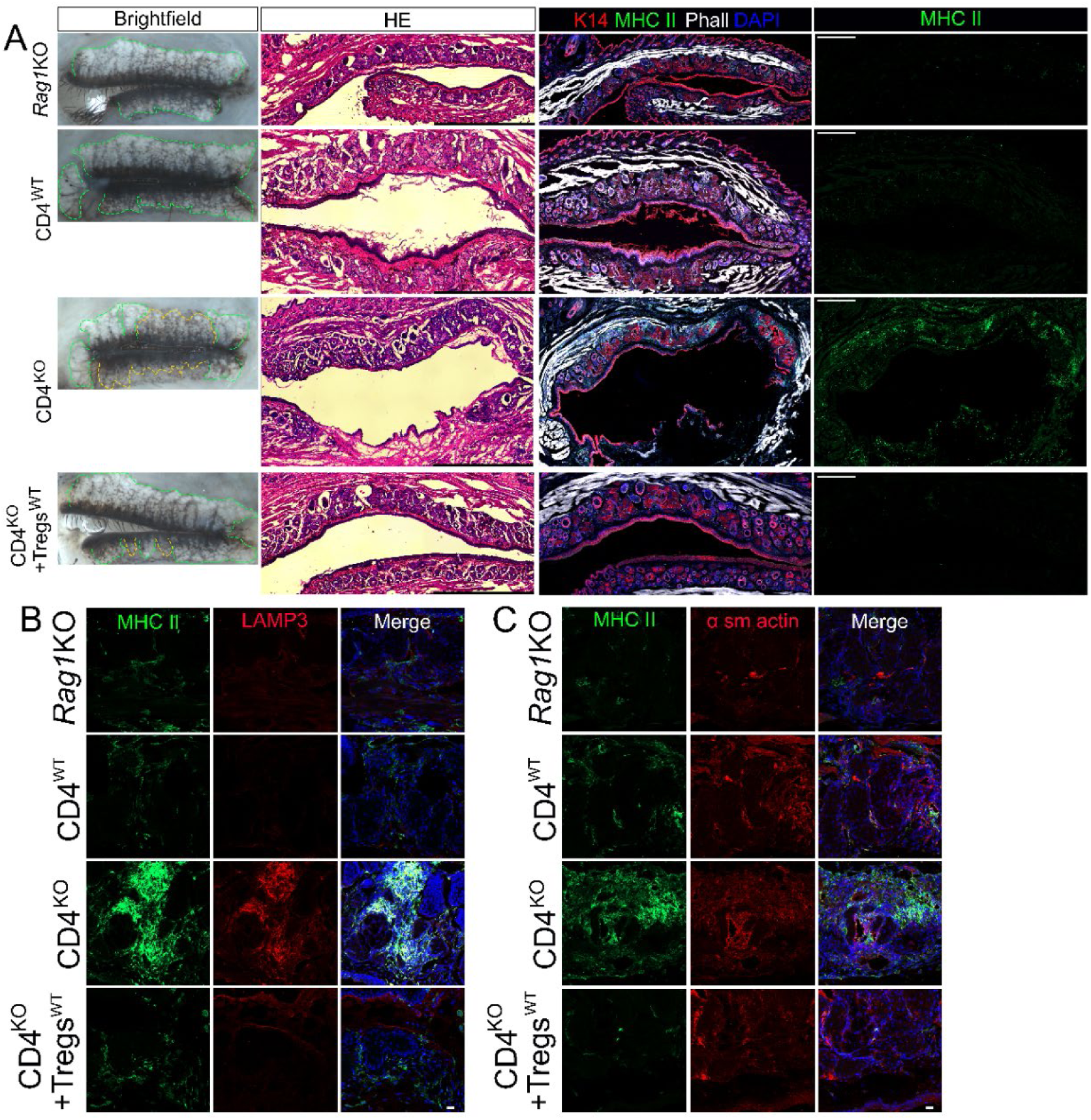
Immunofluorescence identifies exacerbated MHC class II expression in MGs that received only autoimmune CD4^+^ T cells. (**A**) Whole-mounted eyelids imaged under the stereomicroscope with white light, with the outer boundary of the MGs demarcated (first panel). Healthy MGs are demarcated in green, while MGs that are undergoing atrophy are demarcated in yellow. The eyelids were then flat-mounted and processed for cryosectioning, coronal sections obtained, and either stained with H&E (second panel) or immunostained with anti-MHC class II (MHC II - green), anti-Keratin 14 (K14 - red), filamentous actin stained with phalloidin (Phall - white) and nuclei counterstained with DAPI (blue) (third and fourth panels). Images were captured under a stereomicroscope (first panel), an inverted microscope (second panel) using the 10X objective with the spiral mode to image the entire MG area, or a confocal microscope (third and fourth panels) using the 20X objective with the tiling mode to image the entire MG area. The representative images presented in each panel were obtained from the same eyelid sample. The scale bar in the second panel represents 1000 µm, and 500 µm in the third and fourth panel. (**B**) Eyelid cryosections were immunostained with (left) anti-MHC class II (green) anti-LAMP3 (red), and nuclei were counterstained with DAPI (blue) and (right) with anti-MHC II (green using the anti-II-A/I-E antibody), anti-α smooth actin (red), and nuclei were counterstained with DAPI (blue). Scale bar represents 20 µm.

LAMP3, known as lysosomal-associated membrane protein 3 and dendritic cell-lysosomal associated membrane protein, is used as a marker for fully mature dendritic cells(53). MHC class II is primarily expressed by APCs, and dendritic cells are the most potent type of APCs. To investigate the presence of dendritic cells, eyelids were co-stained for MHC II and LAMP3 (Figure 5B). CD4^KO^ recipient mice presented a significant increase in dendritic cell infiltration, whereas a limited number of dendritic cells were identified surrounding the MGs of *Rag1*KO, CD4^WT^ and CD4^KO^+Treg^WT^ recipient mice. Importantly, a significant increase in DCs was noted primarily surrounding atrophic MGs.

### Autoimmune MGD is accompanied by significant fibrosis

To verify whether the increased inflammatory cell infiltration could lead to fibrosis, the tissues were further stained for α smooth muscle actin along with MHC class II, using anti-I-A/I-E, which is a murine-specific MHC class II (Figure 5C). Indeed, in CD4^KO^ recipient mice, areas with increased inflammatory cell infiltration and MHC class II expression exhibited increased α-smooth muscle actin expression, indicating that increased inflammatory cell infiltration coincided with local fibrosis. To better characterize the fibrosis that is induced by autoreactive CD4^+^T cells, the coronal sections of entire eyelids were stained with Masson’s Trichrome staining and hyaluronan (HA). Using Masson’s Trichrome staining, collagen fibers are stained green (Figure 6). Unorganized collagen deposition can be seen in the superior and inferior eyelids of CD4^KO^ recipient mice, however not present in the eyelids of *Rag1*KO or in CD4^WT^ and CD4^KO^+Treg^WT^ recipient eyelids. CD4^KO^ recipient mice presented a significant loss of HA expression surrounding the basal layer of the MG, and an HA-rich fibrotic deposition was observed within the areas where MGs had undergone atrophy (Figure 6B). Our group has previously shown that a net-like distribution of HA surrounding the basal cells is essential for maintaining them in a proliferating state, and that following MG atrophy HA is a major component of the fibrotic deposition that replaces the MGs{Verma, 2025 #9681;Sun, 2018 #9730}.

**Figure 6:**
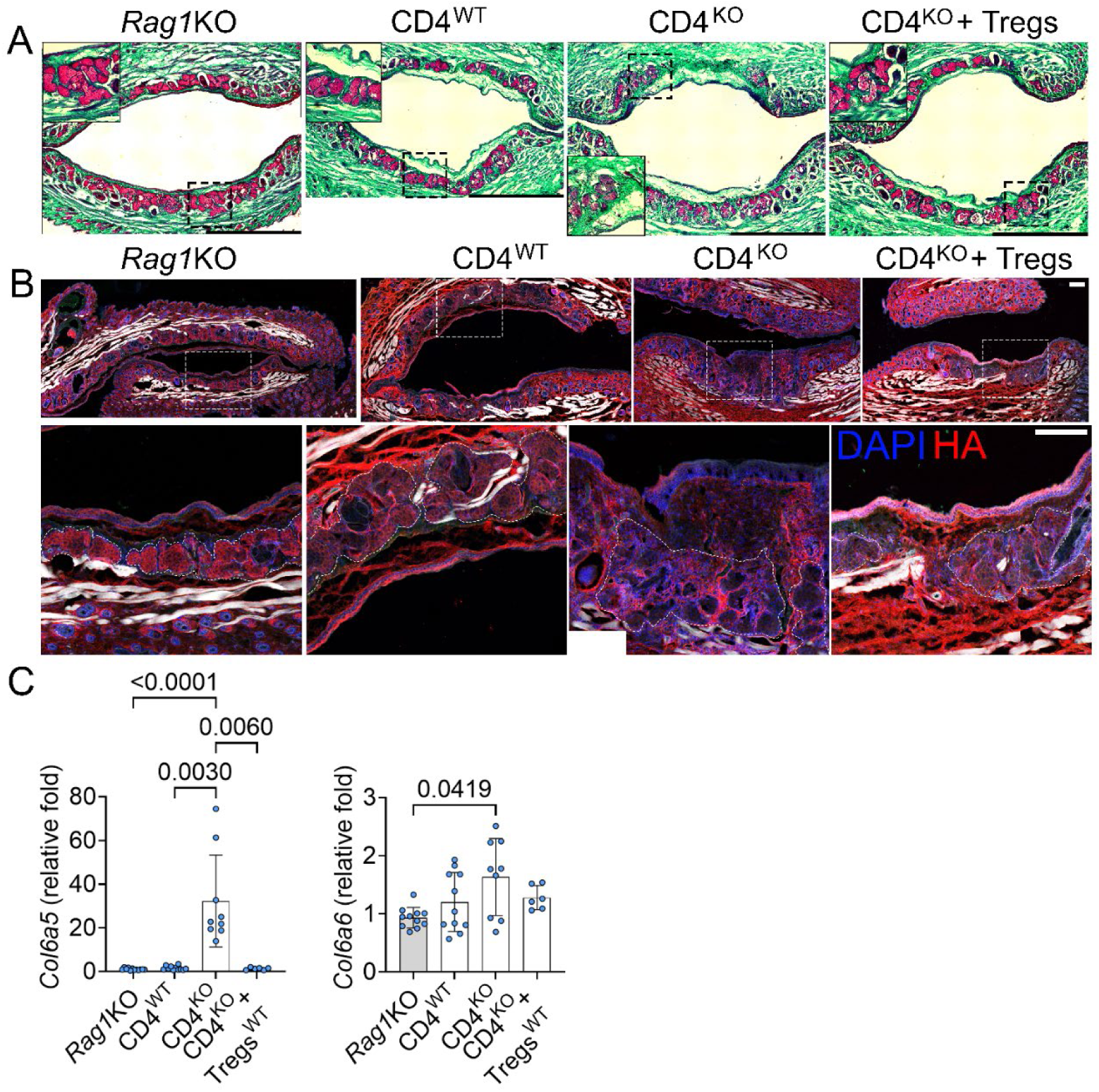
MG fibrosis is evidenced by an increased in collagen deposition and a decreased in hyaluronan expression in the basal layer of MG in CD4^KO^ recipients. (A) Coronal sections of the MG were stained using Masson’s Trichrome. Collagen fibers are observed in green, muscle and cytoplasm appear in red, and nuclei in black. Images were acquired using an inverted microscope with a 20x objective in spiral mode. (B) Eyelid sections were immunostained with anti-HA (red), and nuclei were counterstained with DAPI (blue). Images were captured using a 20x objective in the confocal microscope. C. CD4^KO^ recipients upregulate *Col6a5* gene. Individual Male tarsal plates were collected and subjected to qPCR-based gene expression analysis, which validated *Col6a6* and *Col6a5* involved in fibrosis. P-value as shown. Each dot represents a single male tarsal plate from a different mouse. Mean± SD. One-way ANOVA with Dunn’s multiple comparisons test.

Increased levels of collagen VI have been implicated in fibrosis(54). There are 6 collagen VI genes encoding for different chains(55). Among all collagen VI genes, only *Col6a5* was upregulated in the CD4^KO^/CD4^WT^ (0.84 log2fold change, p-adjusted = 0.01) and CD4^KO^/CD4^KO^+Treg^WT^ comparisons (-1.23 log2fold change, p-adjusted = 0.000203, Supplemental file 1). We validate the expression of *Col6a5* and *Col6a6* using qPCR-specific primers. *Col6a5* and *Col6a6 encode* genes encoding for α5(VI) and α6(VI) collagen VI chains(55). Elevated expression of *Col6a5* was only found in CD4^KO^ recipients (a 30-fold increase compared to naïve *Rag1*KO), while *Col6a6* was only mildly elevated (∼1.7-fold increase). Taken together, our findings indicate that CD4^+^T cell infiltration within the tarsal plate leads to fibrosis in and around the MG.

### A decreased in lipid production is caused by autoimmune MGD

To verify whether the increased inflammatory cell infiltration and fibrosis could affect MG lipid production, the eyelids from *Rag1*KO, CD4^WT^, CD4^KO^, and CD4^KO^+Treg^WT^ recipient mice were processed for ORO staining. CD4^KO^ recipient mice had decreased ORO staining intensity when compared to *Rag1*KO, CD4^WT^, and CD4^KO^+Treg^WT^ recipients, indicating CD4^KO^ mice present an impaired lipid production (Figure 7).

**Figure 7:**
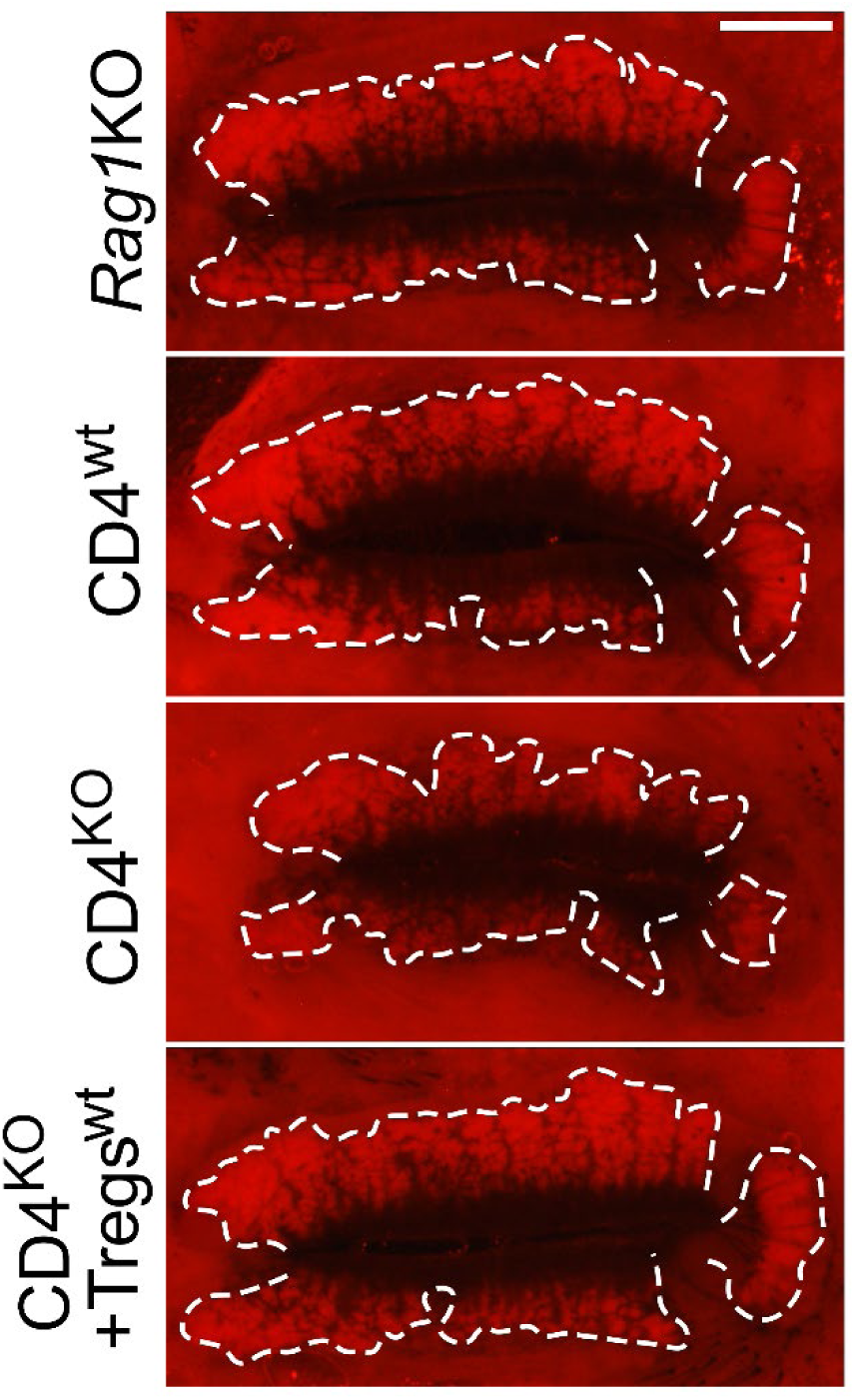
Decreased lipid production in CD4KO recipients. Whole-mounted eyelids were stained with ORO, flat mounted and imaged under a stereomicroscope. Representative digital images are shown. The outer perimeter of the MGs is outlined in white. Scale bar represents 1 mm.

## Discussion

Our results demonstrated for the first time that autoreactive CD4^+^T cells are sufficient to cause MGD in immunodeficient mice. Specifically, autoreactive CD4^+^T cells lead to the accumulation of IFN-γ-producing CD4^+^T cells in the MG periglandular area and upregulation of multiple pathways involved in immune activation, particularly Type II Interferon Signaling which ultimately cause fibrosis, a decrease in lipid production, and MG dropout.

In the clinic, MGD is classified as a chronic and diffuse abnormality of the MGs, that ultimately leads to changes in the quantity and quality of the meibum(4). Changes in the MG area directly affect MG lipid output and are an important metric for assessing MGD. The MG area was quantified in the eyelids of *Rag1*KO, CD4^WT^, CD4^KO^, and CD4^KO^+Treg^WT^ mice, and CD4^KO^ mice presented a decrease in the MG area, although it was more pronounced in the inferior lid. Previous studies have demonstrated that MG atrophy occurs with different incidence and severity in the superior and inferior lids depending on the patient cohort analyzed. For example, MG atrophy is more common in the inferior lid in younger individuals (56), whereas upper MG atrophy is more common in post-menopausal women with acquired nasolacrimal duct obstruction(57). The increased MG atrophy driven by autoimmune CD4^+^T cells, primarily within the inferior lid, remains to be investigated; however, this finding underscores the need to analyze both eyelids in clinical practice. Furthermore, the total number of MGs and the number of enlarged ducts within the eyelids were quantified in *Rag1*KO, CD4^WT^, CD4^KO^, and CD4^KO^+Treg^WT^ mice. A significant decrease in the total number of MGs and a significant increase in the number of enlarged glands within the upper and lower eyelids was observed in the CD4^KO^ mice when compared to *Rag1*KO, CD4^WT^ and CD4^KO^+Treg^WT^ mice, with the difference being more pronounced in the inferior lid. MG plugging is an important early feature of MGD and MG plugging is believed to precede MG atrophy and dropout(31). MG plugging, or obstruction, leads to meibum stasis and backpressure, which eventually causes ductal dilation and MG atrophy {Moreno, 2023 #9710}.

We next characterized the mechanism by which the autoimmune CD4^+^T cells cause MG atrophy. We previously showed that autoreactive CD4^+^T cells isolated from CD25KO mice causes severe LG inflammation and atrophy in young immunodeficient *Rag1*KO recipients(39), while adoptive transfer of naïve C57BL/6 WT CD4^+^T cells does not(58). Further, using this model, the LGs are heavily infiltrated by T cells following the adoptive transfer of autoimmune CD4^+^T cells, leading to severe LG damage within 5 weeks(38, 39). Herein, we found that CD4^+^T cells also infiltrate the periglandular MG area of CD4^KO^ recipient mice. The role of CD4^+^T cells is well established in SjD, with animal models and human exocrine glands and conjunctival biopsies demonstrating that immune infiltrates in these tissues contain a mix of T and B cells(59–61). As we reported in the LG[45], co-adoptive transfer of young WT Tregs with autoreactive CD4^+^T cells prevents MGD damage in young immunodeficient *Rag1*KO recipients. Thus, our results indicate an autoimmune component directly affecting the MG occurs in parallel with LG damage. Furthermore, it suggests that cell therapies targeting Treg function or numbers would be beneficial in SjD.

Th1 (IFN-γ-producing) and Th17 (IL-17-producing) cells have been implicated in dry eye(62, 63). In our results, the infiltration of autoimmune CD4^+^T cells into the periglandular MG area of CD4^KO^ mice dysregulated local immune pathways, including IFN-γ and genes in the IFN-γ pathway. This altered immune milieu led to increased Th1 cell and APC infiltration, in parallel with decreased Th17 cell infiltration into the tarsal plate and MGs of CD4^KO^ mice, compared with all other groups. In an allergic eye disease MG model, which involves immunization of mice with ovalbumin, followed by an OVA topical challenge after 2 weeks, neutrophils and Th17 cells were identified as the major pathogenic immune cells(31). While an increase in neutrophil infiltration in the periglandular area of the MG was observed in *Awat2*KO tarsal plates(64). Our study did not investigate these neutrophils, and this is a promising avenue for further research. In humans, both Th1 and Th17 have been implicated in SjD(65, 66). Genetic deletion experiments showed that deletion of IFN-γ in the CD25KO strain led to delayed dacryoadenitis development while genetic deletion of IL-17 in the same parental strain led to accelerated LG inflammation(67). However, Th1/Th17/Th2 cells can influence the differentiation of the others(68) so it is not surprising that one subset may dominate in certain models. In a clinical setting, IFN-γ has been associated with SjD, for example, an increased “IFN” signature is present in the sera and conjunctiva of SjD(69–73). Increased levels of IFN-γ have also been shown to mediate tissue damage, with a subconjunctival injection of IFN-γ decreasing goblet cell density in mice using an experimental dry eye model(62). Given that CD4^KO^+Treg^WT^ recipients showed a decrease in IFN-γ and a modest increase in Th17 cells, this could suggest that, the Th1 cells are the main pathogenic cells in our model; however, future studies are needed to verify this.

IFN-γ regulates the expression of *Ciita*, the master regulator for MHC class II expression. High levels of IFN-γ can lead to MHC class II expression in certain non-APC populations, such as fibroblasts, epithelial cells, and endothelial cells(74–76). By bulk RNA sequencing, qPCR, and immunostaining, MHC class II was identified as a key molecule upregulated within the periglandular MG area in CD4^KO^. Importantly, in the CD4^KO^ recipient mice, MHC class II expression was not limited to APCs; with epidermal keratinocytes and stromal fibroblasts also expressing MHC class II. MHC class II is a cell surface protein that presents peptide fragments to CD4^+^ T cells, which is a key step in the adaptive immune response. In healthy tissues, the expression of MHC class II is restricted to APCs(77). However, in certain pathological contexts, MHC class II expression can be induced in resident cells, such as during autoimmune disease(76, 78). The induction of MHC class II expression by non-immune cells can lead to an exacerbated immune response and the immune recognition of self-antigens(79, 80). The expression of MHC class II in non-APCs occurs in various autoimmune diseases, such as SjD, Hashimoto thyroiditis, rheumatoid arthritis, and multiple sclerosis, and in certain chronic infections or following tissue injury(81).

Our results show an increased co-expression of LAMP3 and MHC II. LAMP3 was originally described as a marker of dendritic cells, but epithelial expression has also been reported. LAMP3 overexpression in mice results in a clinical phenotype of SjD with oral features (LG not evaluated)(82). Additionally, a clinical study evidenced an association of LAMP3 with autoantibodies in SjD(83). Our study investigating immune pathways in the conjunctiva of SjD patients demonstrated increased LAMP3 mRNA levels compared to controls(69). Together, these observations link LAMP3 overexpression to sustained immune activation and chronic inflammation in SjD. Chronic inflammation activates fibroblasts and alters extracellular matrix deposition, resulting in fibrosis (84);Shapouri-Moghaddam, 2018 #8641}. In turn, fibrosis impairs tissue function, which is particularly relevant in MG, as it can cause narrowing or obstruction of the ducts(85). Furthermore, the changes in the biomechanical properties of the tissue following fibrosis could also affect MG function. We also investigated whether autoimmune CD4^+^T cells cause fibrosis within and around the MG. Increased α-smooth muscle actin expression was detected in the MG periglandular area in CD4^KO^ mice compared to all other groups, indicating an increased number of myofibroblasts surrounding the MG at 5 weeks after adoptive transfer, suggesting that the increased inflammatory cell infiltration elicits local fibrosis. Eyelids were stained with Masson’s Trichrome staining to study the distribution and organization of collagen within the tarsal plate. Unorganized collagen deposition was observed in both the superior and inferior eyelids of CD4^KO^ recipient mice; however not observed in *Rag1*KO, CD4^WT^, and CD4^KO^+Treg^WT^ mice. We have previously shown HA is organized in an intricate net-like distribution surrounding the basal cells of the MGs, and as cable-like structures in the tarsal plate. In age-related MGD, there is a loss of the net-like distribution of HA surrounding the basal layer of the MG that precedes MG atrophy(25). Following MG atrophy, these glands are replaced by HA organized as a fibrotic deposition (25). Therefore, we also investigated HA expression in the tarsal plate and surrounding the MG in our model. CD4^KO^ recipient mice presented a significant loss of HA expression surrounding the basal cells of the MG following the adoptive transfer of autoimmune CD4^+^T cells, while the other groups did not. Additionally, HA organized in a fibrotic distribution was observed throughout the tarsal plate in CD4^KO^ recipient mice, further supporting the notion that these mice exhibit a fibrotic process in both eyelids. The increase in MHC class II expression and fibrosis was observed in both the superior and inferior eyelids; however, MG atrophy was mostly evident in the inferior lid. Future studies should investigate why the inferior lid was more susceptible to MG atrophy than the superior eyelid in this model, despite both showing pathological immune changes.

To date, various animal models have been developed to study MGD(31, 86–88). In models that lead to meibomian stasis, such as alkali burn or eyelid margin cauterization, and in models of age-related MGD, MG atrophy occurs primarily in acini furthest from the eyelid opening, leading to gland shortening(25). In our model of autoimmune MGD, MG atrophy was observed throughout the gland, with notable narrowing and shortening. Ghost glands were also evident in, and the presence of ghost glands shows an irreversible structural degeneration of the MG. Their appearance in this autoimmune MGD model further demonstrates that immune-driven injury accelerates the progression of MGD. Future studies should be dedicated to characterizing the dynamics of MG atrophy among the different models of MGD to aid our understanding of how different pathological mechanism affect MG homeostasis.

This study found that, as with the LG and salivary glands, autoimmune CD4^+^T cells also infiltrate the periglandular MG area, leading to progressive loss of MG tissue. Although both the LG and MG exhibit similar pathological findings, disease onset and rate of progression are significantly faster in the LG than in the MG. In fact, by 4 to 5 weeks, all mice present severe loss of LG tissue (38, 39), whereas the MG presents early signs of MGD at this same time point. A limitation of this study was that we did not extend our evaluation past 5 weeks of post-adoptive transfer.

The dichotomy of dry eye classification as “ADDE” or “evaporative dry eye” is widely used clinically. However, mixed cases (also called hybrids) have been described(4). Taken together, our results show for the first time that autoreactive CD25KO CD4^+^T cells cause autoimmune MGD in addition to dacryoadenitis. This finding has implications for the clinical management of SjD and underscores the importance of examining dry eye globally rather than being constrained to the binary classification of ADDE or evaporative dry eye. Furthermore, our results suggest that cell-based therapies that augment Treg suppressive function would also prevent autoimmune MDG in these patients.

## Acknowledgments

We gratefully acknowledge Leiqi Zhang’s contribution to the management of the mouse colonies.

## Supplemental Figures

**Supplemental Figure 1:**
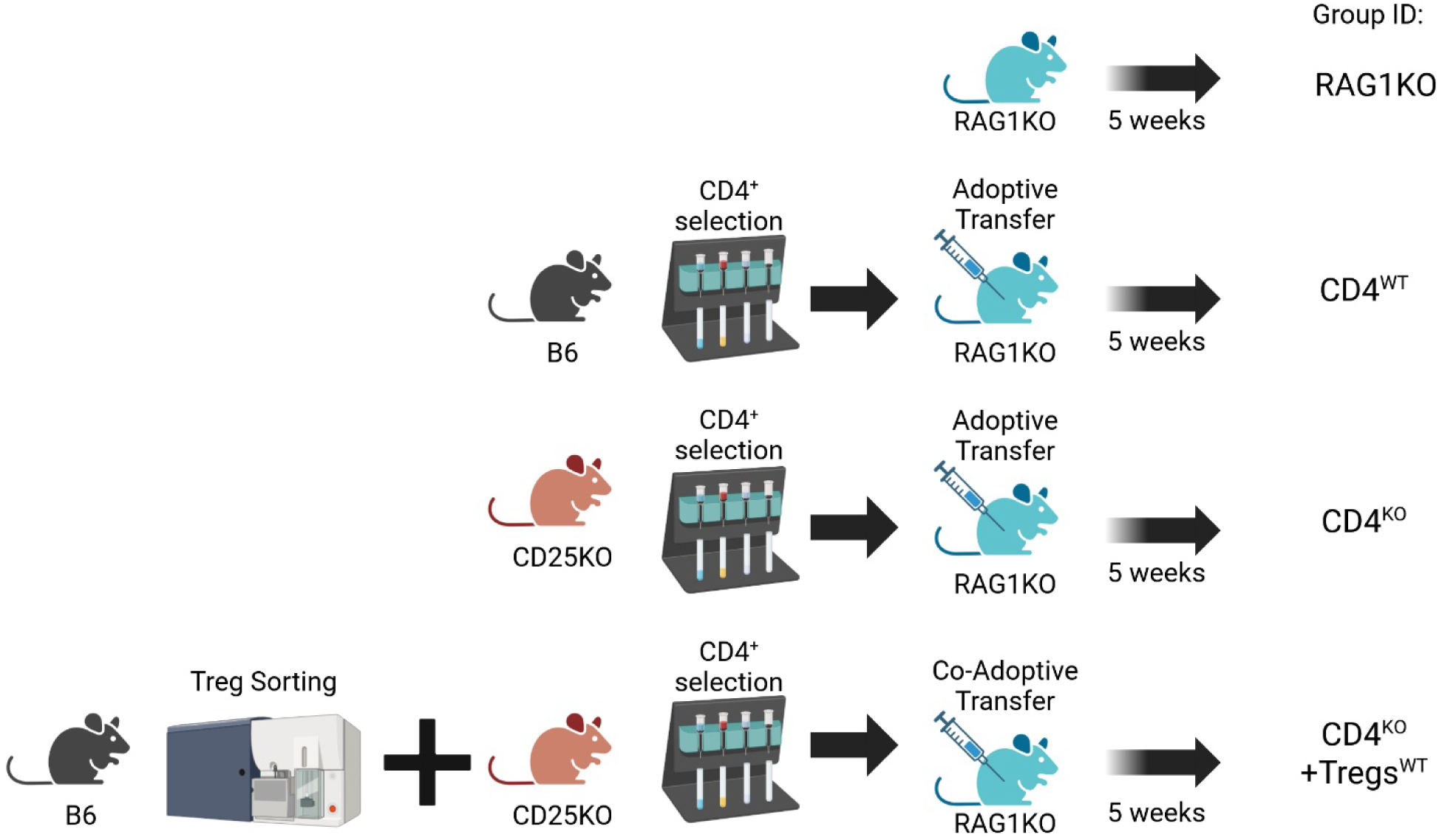
Adoptive transfer schematic. Cells were isolated as described in the methods. B6 = C57BL/6J, CD25^+/+^.

**Supplementary Figure 2:**
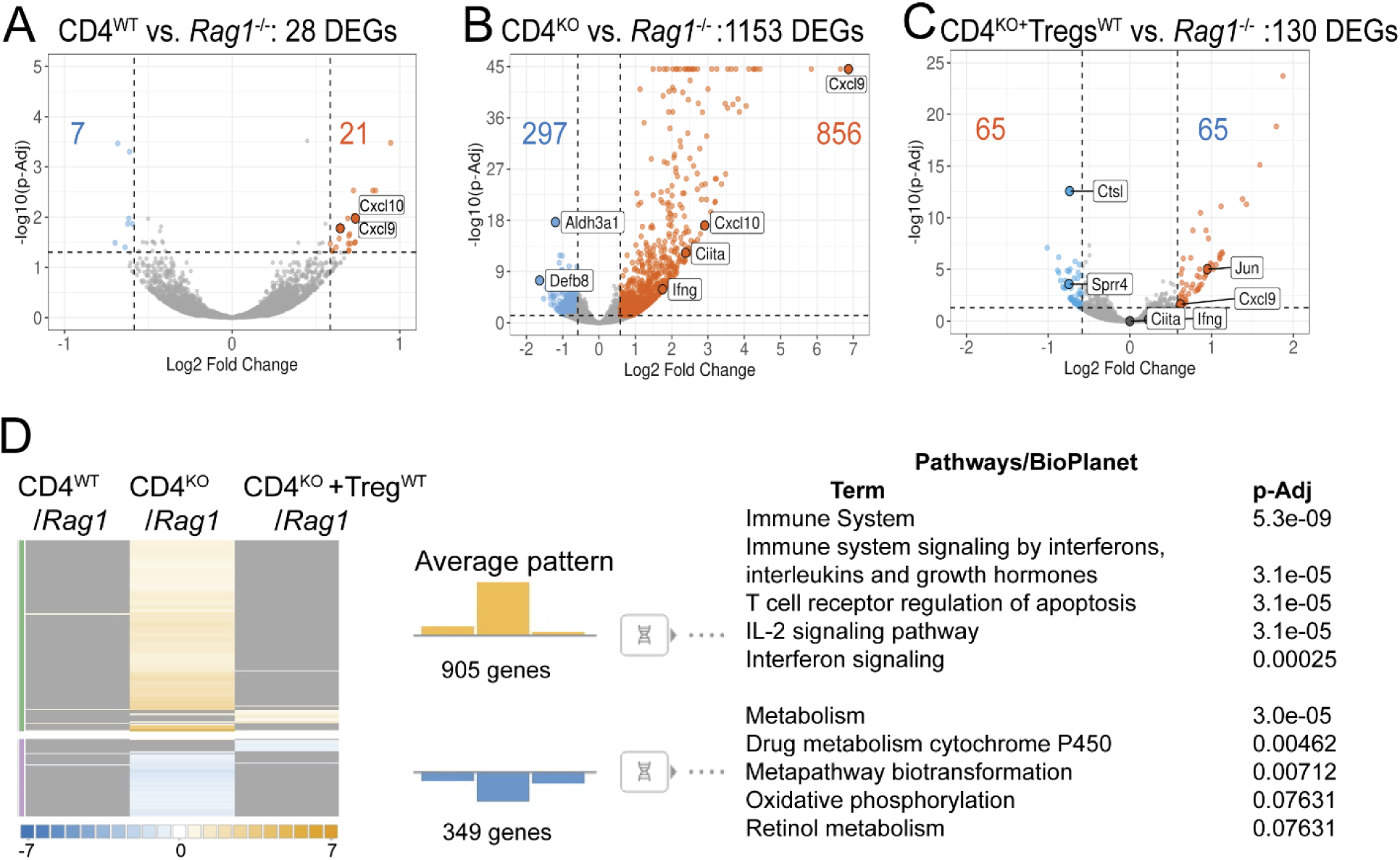
The expression of inflammatory pathways within the MG changes based on source of T cells adoptively transferred. Tarsal plates were collected after 5 weeks post-transfer and lysed. Total RNA was subjected to bulk-RNA sequencing. The *Rag1*KO tarsal plates were used as calibrators. **A-B** Volcano plots demonstrate the magnitude of change comparing CD4^WT^ to Rag1KO (**A**), CD4^KO^ to *Rag1*KO (**B**) and CD4^KO^+Tregs^WT^ to *Rag1*KO (**C**) tarsal plates. **D**. Metanalysis indicating unique upregulated (yellow)/downregulated (blue) pathways.

