## Supplemental Figures for "Autoimmune CD4^+^T cells Cause Meibomian Gland Dysfunction"

**Supplemental Figure 1**

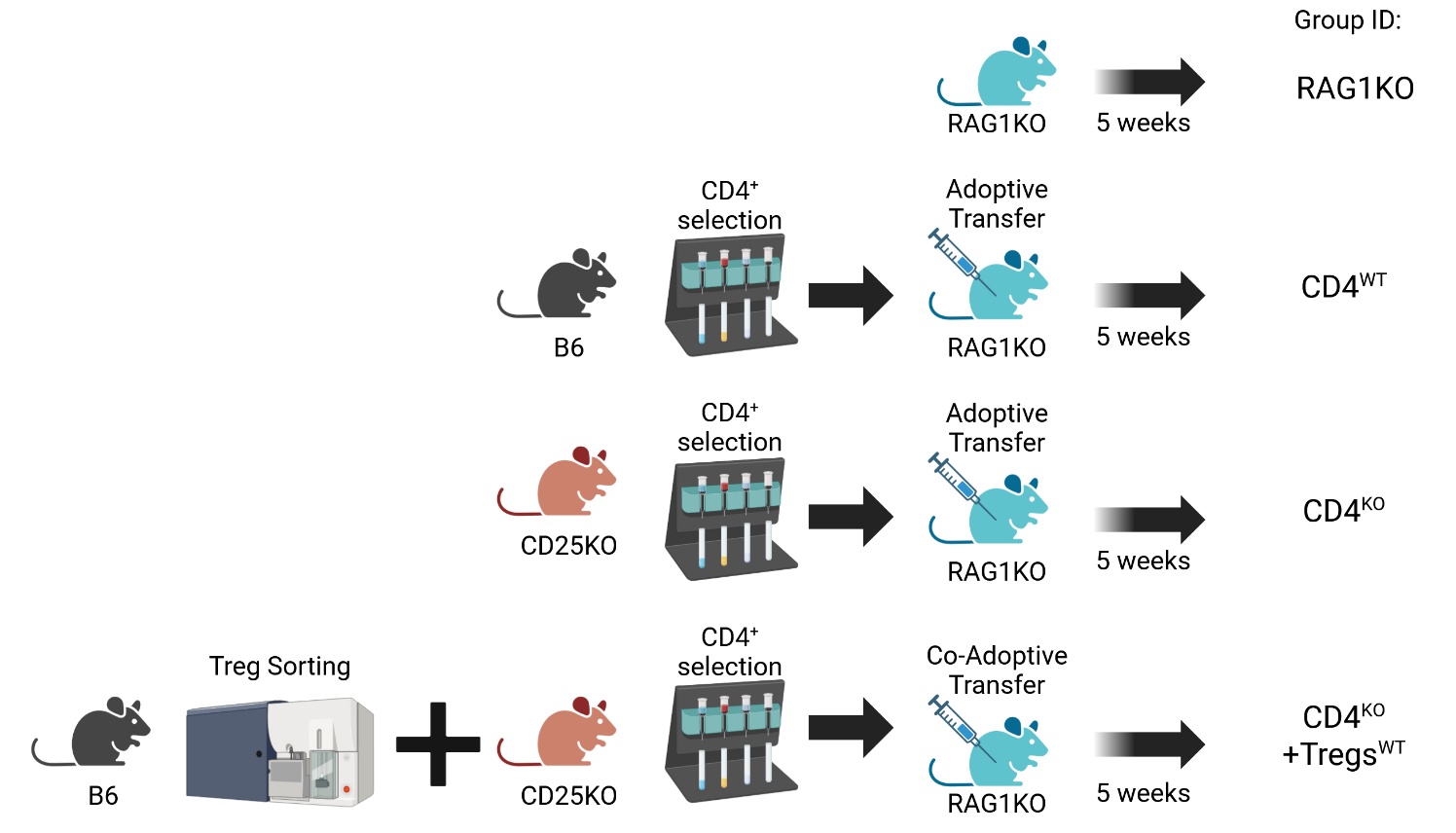

**Supplemental Figure 1: Adoptive transfer schematic.** Cells were isolated as described in the methods. B6 = C57BL/6J, CD25^+/+^.

**Supplemental Figure 2**

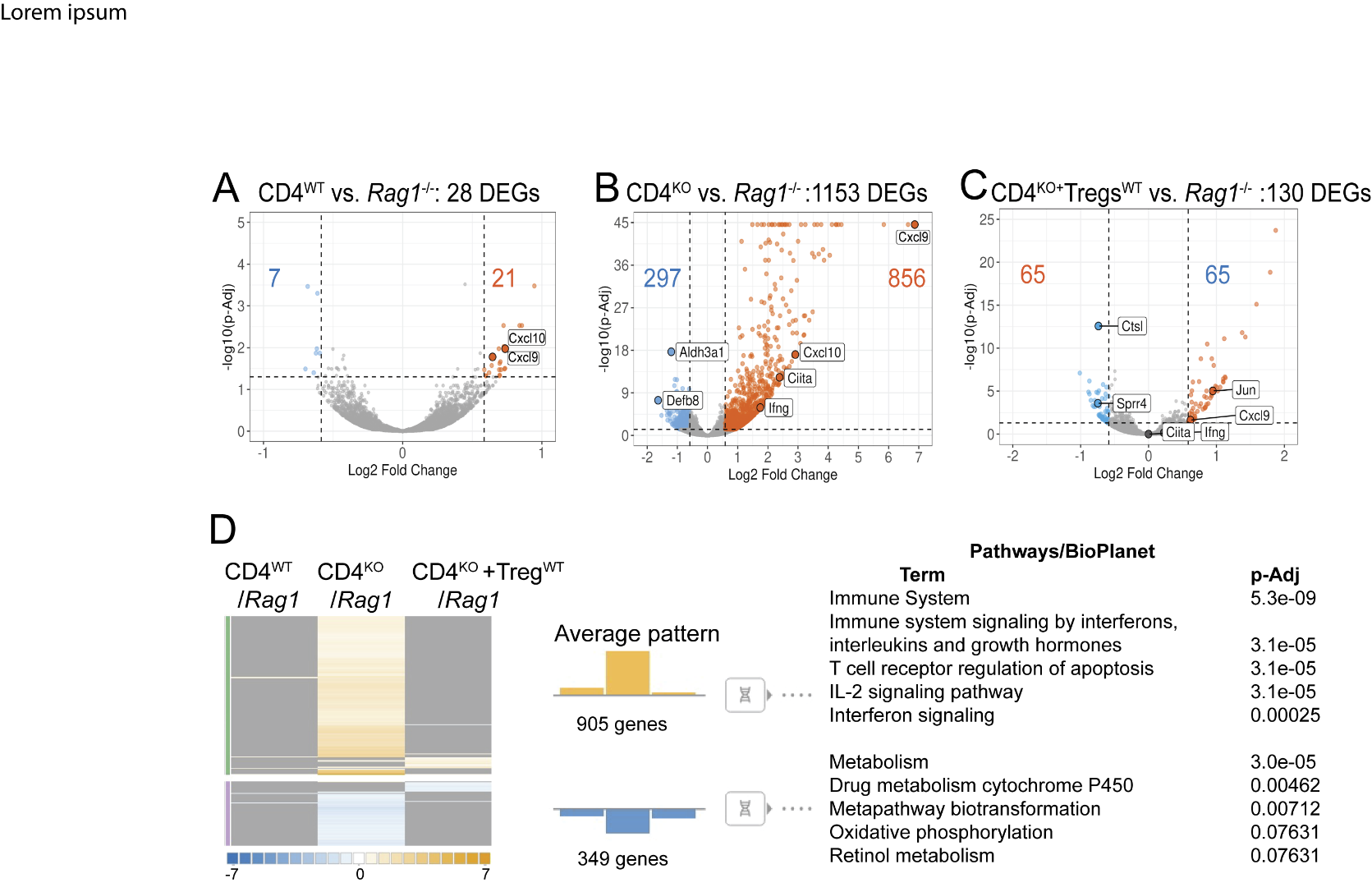

**Supplementary Figure 2:** The expression of inflammatory pathways within the MG changes based on source of T cells adoptively transferred. Tarsal plates were collected after 5 weeks post-transfer and lysed. Total RNA was subjected to bulk-RNA sequencing. The *Rag1*KO tarsal plates were used as calibrators. **A-B** Volcano plots demonstrate the magnitude of change comparing CD4^WT^ to Rag1KO (**A**), CD4^KO^ to *Rag1*KO (**B**) and CD4^KO^+Tregs^WT^ to *Rag1*KO (**C**) tarsal plates. **D**. Metanalysis indicating unique upregulated (yellow)/downregulated (blue) pathways.
